# 11β-Hydroxysteroid Dehydrogenase Type 1 inhibition in Idiopathic Intracranial Hypertension: a double-blind randomized controlled trial

**DOI:** 10.1101/648410

**Authors:** Keira Markey, James Mitchell, Hannah Botfield, Ryan S Ottridge, Tim Matthews, Anita Krishnan, Rebecca Woolley, Connar Westgate, Andreas Yiangou, Pushkar Shah, Caroline Rick, Natalie Ives, Angela E Taylor, Lorna C Gilligan, Carl Jenkinson, Wiebke Arlt, William Scotton, Rebecca Fairclough, Rishi Singhal, Paul M Stewart, Jeremy W Tomlinson, Gareth G Lavery, Susan P Mollan, Alexandra J Sinclair

## Abstract

Treatment options for idiopathic intracranial hypertension are limited. The enzyme 11β-hydroxysteroid dehydrogenase type 1 has been implicated in regulating cerebrospinal fluid secretion, and its activity is associated with alterations in intracranial pressure in idiopathic intracranial hypertension. We assessed therapeutic efficacy, safety and tolerability, and investigate indicators of *in vivo* efficacy of the 11β-hydroxysteroid dehydrogenase type 1 inhibitor AZD4017 compared to placebo in idiopathic intracranial hypertension. A multicenter, UK, 16-week phase II randomized, double-blind, placebo-controlled trial of 12-weeks treatment with AZD4017 or placebo was conducted. Women aged 18 to 55 years with active idiopathic intracranial hypertension (>25cmH_2_O lumbar puncture opening pressure and active papilledema) were included. Participants received 400mg twice daily of oral AZD4017 compared to matching placebo over 12-weeks. The outcome measures were initial efficacy, safety and tolerability. The primary clinical outcome was lumbar puncture opening pressure at 12 weeks analysed by intention-to-treat. Secondary clinical outcomes were symptoms, visual function, papilledema, headache and anthropological measures. *In vivo* efficacy was evaluated in the central nervous system and systemically. 31 subjects (mean age 31.2 (SD=6.9) years and BMI 39.2 (SD=12.6) kg/m^2^) were randomized to AZD4017 (n=17) or placebo (n=14). At 12 weeks, lumbar puncture pressure was lower in the AZD4017 group (29.7 cmH_2_O) compared with placebo (31.3 cmH_2_O), but the difference between groups was not statistically significant (mean difference: −2.8, 95% confidence interval: −7.1-1.5; p=0.2). An exploratory analysis assessing mean change in lumbar puncture pressure within each group found a significant decrease in the AZD4017 group (mean change: −4.3 cmH_2_O (SD=5.7); *p*=0.009) but not in the placebo group (mean change: −0.3 cmH_2_O (SD=5.9); p=0.8). AZD4017 was safe, with no withdrawals related to adverse effects. Nine transient drug-related adverse events were reported. One serious adverse event occurred in the placebo group (deterioration requiring shunt surgery). *In vivo* biomarkers of 11β-hydroxysteroid dehydrogenase type 1 activity (urinary glucocorticoid metabolites, hepatic prednisolone generation and CSF cortisone to cortisol ratios) demonstrated significant enzyme inhibition. This is the first phase 2 randomized controlled trial in idiopathic intracranial hypertension evaluating a novel therapeutic target. AZD4017 was safe, well-tolerated and inhibited 11β-hydroxysteroid dehydrogenase type 1 activity *in vivo*. Possible clinical benefits were noted in this small cohort. A longer, larger study would now be of interest.

## Introduction

Idiopathic intracranial hypertension (IIH) is a debilitating condition characterized by raised intracranial pressure (ICP), papilledema with risk of permanent visual loss,(Mollan *et al.*, 2018b) and chronic headaches which reduce quality of life.(Mulla *et al.*, 2015) IIH predominately affects obese women between the ages of 25-36 years with a distinct androgen excess signature recently identified.(Daniels *et al.*, 2007; Markey *et al.*, 2016; O’Reilly *et al.*, 2019) Incidence is increasing in line with escalating worldwide obesity rates.(Mollan *et al.*, 2018a)

Surgical treatment is recommended when vision rapidly declines,(Mollan *et al.*, 2014; Mollan *et al.*, 2018c) but the majority of patients (93%) are managed conservatively.(Hoffmann *et al.*, 2018; Mollan *et al.*, 2018a; Mollan *et al.*, 2018b) Dietary interventions are an effective treatment,(Sinclair *et al.*, 2010a) however, meaningful and sustained weight loss is difficult to achieve.(Colquitt *et al.*, 2014; Manfield *et al.*, 2017) Pharmacotherapy in IIH is limited,(Piper *et al.*, 2015) with only two previous randomized controlled trials (RCTs) in IIH previously reported, both evaluating acetazolamide.(Ball *et al.*, 2011; Committee *et al.*, 2014) New treatment options are therefore urgently required.(Mollan *et al.*, 2018b)

We have previously demonstrated that the enzyme 11β-hydroxysteroid dehydrogenase type 1 (11β-HSD1) is expressed and active in the choroid plexus (CP) to amplify cortisol availability and acts to regulate cerebrospinal fluid (CSF) production.(Gathercole *et al.*, 2013)’(Sinclair *et al.*, 2007; Sinclair *et al.*, 2010c) In patients with IIH, resolution of disease (reduced ICP, improvements in papilledema and headaches) was associated with reduced 11β-HSD1 activity,(Sinclair *et al.*, 2010a; Sinclair *et al.*, 2010c) with a study suggesting that inhibition of 11β-HSD1 with a non-selective inhibitor lowered intraocular pressure.(Rauz *et al.*, 2003) Importantly, 11β-HSD1 expression and activity is dysregulated in obesity.(Sandeep *et al.*, 2005) (Wake and Walker, 2004)

Selective inhibitors of 11β-HSD1 have been developed as treatments for obesity, hepatic steatosis, metabolic syndrome and type 2 diabetes.(Boyle, 2008; Stefan *et al.*, 2014) Based on these data, 11β-HSD1 could represent a therapeutic target for lowering CSF pressure. AZD4017 is a highly selective, fully reversible, competitive 11β-HSD1 inhibitor. It has been tested over short time intervals in healthy males (9 days), and abdominally obese subjects (10 days), and found to be safe and tolerable.(AstraZeneca, 2000-[01 March 2017]-a, b, c, d, 2000-[01 March 2017].) The ability of AZD4017 to penetrate the blood-brain-barrier (BBB) is not established; however, the CP lies outside the BBB and consequently can be targeted directly following oral administration.(Davson, 1966; Eftekhari *et al.*, 2015)

We hypothesised that inhibition of 11β-HSD1 could be therapeutically beneficial in IIH. To test this we conducted a multicenter phase 2 double-blind, placebo-controlled RCT in IIH using the selective 11β-HSD1 inhibitor AZD4017, aiming to assess therapeutic efficacy, safety and tolerability, and investigate *in vivo* systemic and central nervous system efficacy.

## Methods

### Study Conduct

The study was conducted from March 2014 to December 2016 in three UK hospitals. The National Research Ethics Committee York and Humber-Leeds West gave ethical approval (13/YH/0366). All patients provided written informed consent in accordance with the delaration of Helsinki. Detailed clinical trial methodology has been published.(Markey *et al.*, 2017a)

### Study Population

Women (18-55 years) were eligible if they had a clinical diagnosis of active IIH meeting the updated, modified Dandy criteria (ICP>25cmH_2_O and active papilledema) and normal brain imaging (including magnetic resonance venography or CT with venography) at recruitment (for detailed eligibility criteria see Suppl. Table 1).(Friedman *et al.*, 2013; Markey *et al.*, 2017b)

### Study design

A 16-week phase II, double-blind placebo-controlled RCT with a 12-week dosing duration and 4-week follow-up off drug.

### Randomization and Blinding

Participants were allocated to either drug or placebo. Patients were allocated a trial number randomly by phone, using block-of-6 randomization. Participants and investigators were masked to treatment allocation during the trial.

### Intervention

An oral selective 11β-HSD1 inhibitor, AZD4017, at 400mg twice daily for 12-weeks, compared to a matched placebo. Trial dosing was added to existing therapy for IIH, other drugs were maintained at a fixed dose throughout the study.

### Assessments

Participants completed follow-up assessments at 1, 2, 3, 4, 6, 8, 10, 12 and 16 weeks (Figure 1A).

**Figure 1:**
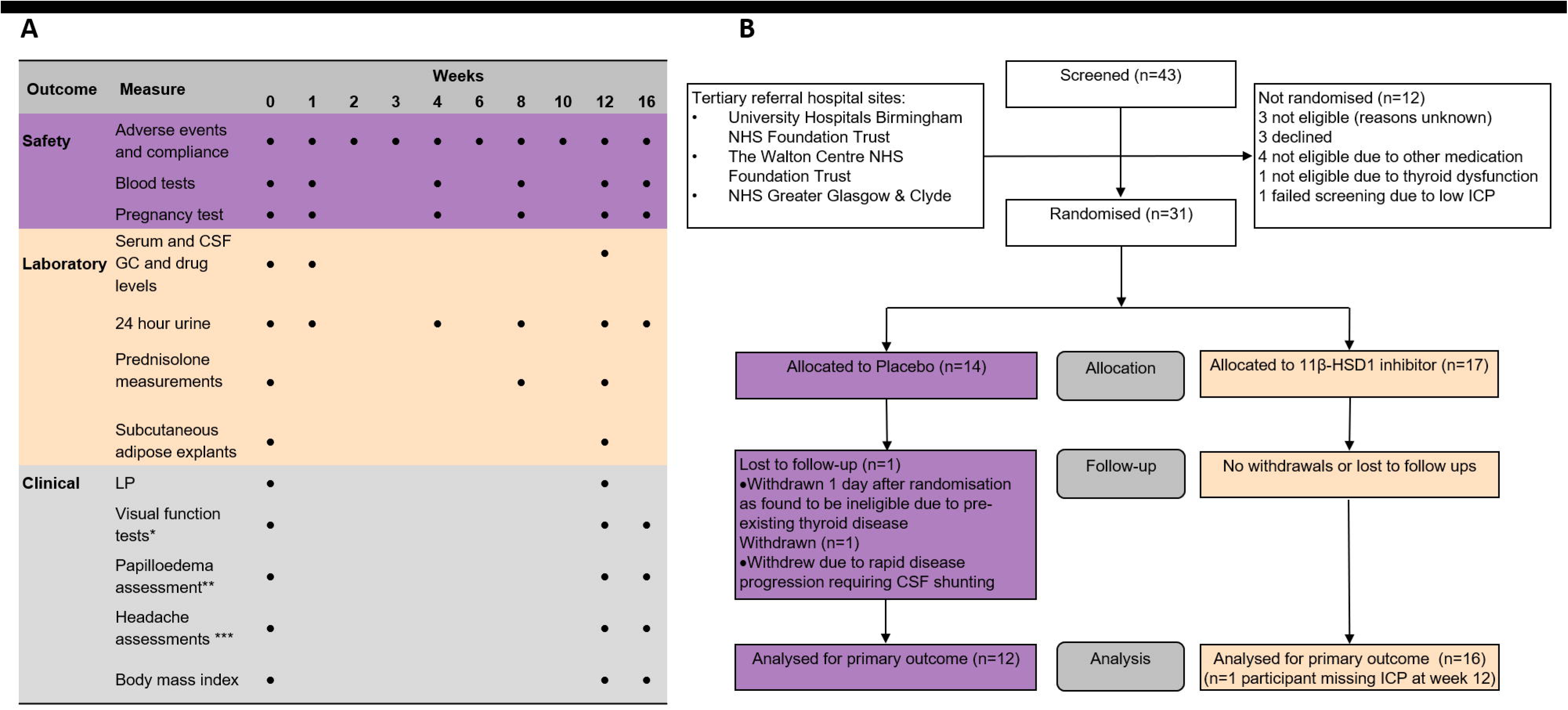
Participant visits (A) and CONSORT diagram (B). GC, glucocorticoids, CSF, cerebrospinal fluid, ICP, intracranial pressure.

#### Clinical assessments

The primary outcome for clinical efficacy was the difference in ICP between AZD4017 and placebo, as measured by LP at 12 weeks. Secondary outcomes included: IIH symptoms, visual function (visual acuity (VA) measured using LogMAR (log of the minimum angle of resolution), perimetric mean deviation (PMD) using Humphrey 24-2 central threshold automated perimetry and contrast sensitivity assessed by MARs charts (Mars Perceptrix, USA)), papilledema, headache associated disability and anthropological measures. Papilledema was evaluated using spectral domain optical coherence tomography (OCT; Spectralis, Heidelberg Engineering) to quantify the peripapillary retinal nerve fibre layer (RNFL) average and maximal values. Papilledema was graded from fundus photographs by three masked neuro-ophthalmologists using the Frisén classification (0 denotes no papilledema to grade 5 severest papilledema).(Frisen, 1982) Headache was evaluated through the headache impact test-6 disability questionnaire (HIT-6), headache severity (verbal rating score 0 to 10), frequency (days per month), duration and analgesic use (days per month).(Bayliss *et al.*, 2003) Pill counting at each visit determined drug compliance.

In the original grant application and early versions of the protocol, the primary outcome measure was stated as the change in ICP between baseline and 12 weeks. Following adoption of the study by the Birmingham Clinical Trials Unit, the primary outcome was changed to ICP at 12 weeks, with adjustment for baseline ICP in the analysis. This change was made blind to any data analysis.

#### Safety and Tolerability

Adverse events and safety bloods were monitored (timeline Figure 1A) including renal function (urea, creatinine and electrolytes), liver function (aspartate transaminase, alanine transferase, bilirubin, alkaline phosphatase, gamma-glutamyl transferase), thyroid function (thyroid stimulating hormone, free thyroxine) and creatine kinase. Hypothalamic pituitary adrenal (HPA) axis activity was monitored (cortisol, adrenocorticotropic hormone (ACTH), dehydroepiandrosterone sulfate (DHEAS), testosterone, androstenedione, follicle stimulating hormone, luteinizing hormone, estradiol and progesterone).

#### Glucocorticoid and AZD4017 blood and CSF levels

Samples were collected and stored at −80°C. Cortisol and cortisone levels in serum and CSF were measured by liquid chromatography-tandem mass spectrometry (LC-MS/MS) at the University of Birmingham, as previously described.(Hassan-Smith *et al.*, 2015; Long *et al.*, 2016) Plasma and CSF AZD4017 levels were quantified by an external laboratory (Alderley Analytical, Knutsford, UK).

#### In vivo systemic 11β-Hydroxysteroid Dehydrogenase activity

Global 11β-HSD1 activity was evaluated through quantification of 24 hour urinary glucocorticoid metabolites, by LC-MS/MS.(Sagmeister *et al.*, 2018) 11β-HSD1 activity was inferred from the ratio of (5α-tetrahydrocortisol + tetrahydrocortisol):tetrahydrocortisone ((5αTHF+THF):THE) alongside a stable ratio of total urinary cortisol (F): total urinary cortisone (E) reflecting 11β-HSD2 activity.^(Tomlinson and Stewart, 2001)^

#### In vivo hepatic 11β-Hydroxysteroid Dehydrogenase activity

Inhibition of hepatic 11β-HSD1 activity was informed by measuring first-pass metabolism of 10mg of oral prednisone to prednisolone. Serum prednisone and prednisolone were measured every 20 minutes over 4 hours using LC-MS/MS.(Richards *et al.*, 2012; Hassan-Smith *et al.*, 2015)’

#### Ex vivo adipose 11β-Hydroxysteroid Dehydrogenase activity

Subcutaneous adipose biopsies (100-150 mg distributed to triplicate experiment) incubated in media (Dulbecco’s Modified Eagle Medium/Nutrient Mixture F-12 (ThermoFisher, Rugby, UK) at room temperature with 100 nM cortisone (Sigma-Aldrich, Dorset, UK), with three media controls (without adipose) for 24 hours. Steroid conversion was quantified using LC-MS/MS.(Juhlen *et al.*, 2015; Mooij *et al.*, 2015)

#### In vitro adipose 11β-Hydroxysteroid Dehydrogenase inhibition by AZD4017

Subcutaneous and omental adipose explants (1-2g in triplicate) were obtained from IIH patients undergoing bariatric surgery. Samples were incubated with either 2000nM, 200nM or 20nM of AZD4017 and 100nM of cortisone alongside three controls (without AZD4017) for 24 hours. Steroid conversion was quantified using LC-MS/MS.(Juhlen *et al.*, 2015; Mooij *et al.*, 2015)

### Statistical Analysis

Analysis of the clinical data was based on the full analysis set according to the statistical analysis plan (Supplemental document). Analysis was conducted using intention-to-treat with data from all available randomized participants used. The primary comparison was between AZD4017 versus placebo at 12 weeks. The majority of data was continuous, so groups were compared using linear regression models with baseline measurements included as a covariate in the model. IIH symptom data was binary and was analyzed using log-binomial models with baseline symptom included as a covariate in the model. The primary analysis of visual data included data from both eyes, using a linear mixed model with participant included as a random effect. We also analyzed data from the most affected eye at baseline as defined by PMD.(Friedman *et al.*, 2014) Statistical significance was set at p<0.05, with no adjustment for multiple comparisons made. Clinical data was analyzed using SAS (version 9.4) and STATA (version 14).

Analysis of laboratory data was performed using SPSS (version 24, IBM, New York, USA). An unpaired t-test was used for normally distributed data (Mann-Whitney U test for non-parametric data). For related groups, either the paired t-test or Wilcox signed-rank test was used for parametric and non-parametric data respectively. We reported means and standard deviations (medians and ranges for non-parametric data).

#### Sample Size

To detect a difference between groups of 14% in ICP (assuming a standard deviation of 10% for ICP) with 90% power and two-sided alpha=0.05, required 12 participants per group. Allowing for 20% drop-out, we aimed to recruit 30 participants.

#### Data availability

The trial is registered at Clinicaltrials.gov NCT02017444; European Clinical Trials Database (EudraCT Number: 2013-003643-31). The data that support the findings of this study are available from the corresponding author, upon reasonable request.

## Results

31 participants were recruited; 17 were randomized to AZD4017 and 14 to placebo (Figure 1B). Baseline characteristics demonstrate a cohort of IIH patients with active disease were recruited (Table 1).

**Table 1:**
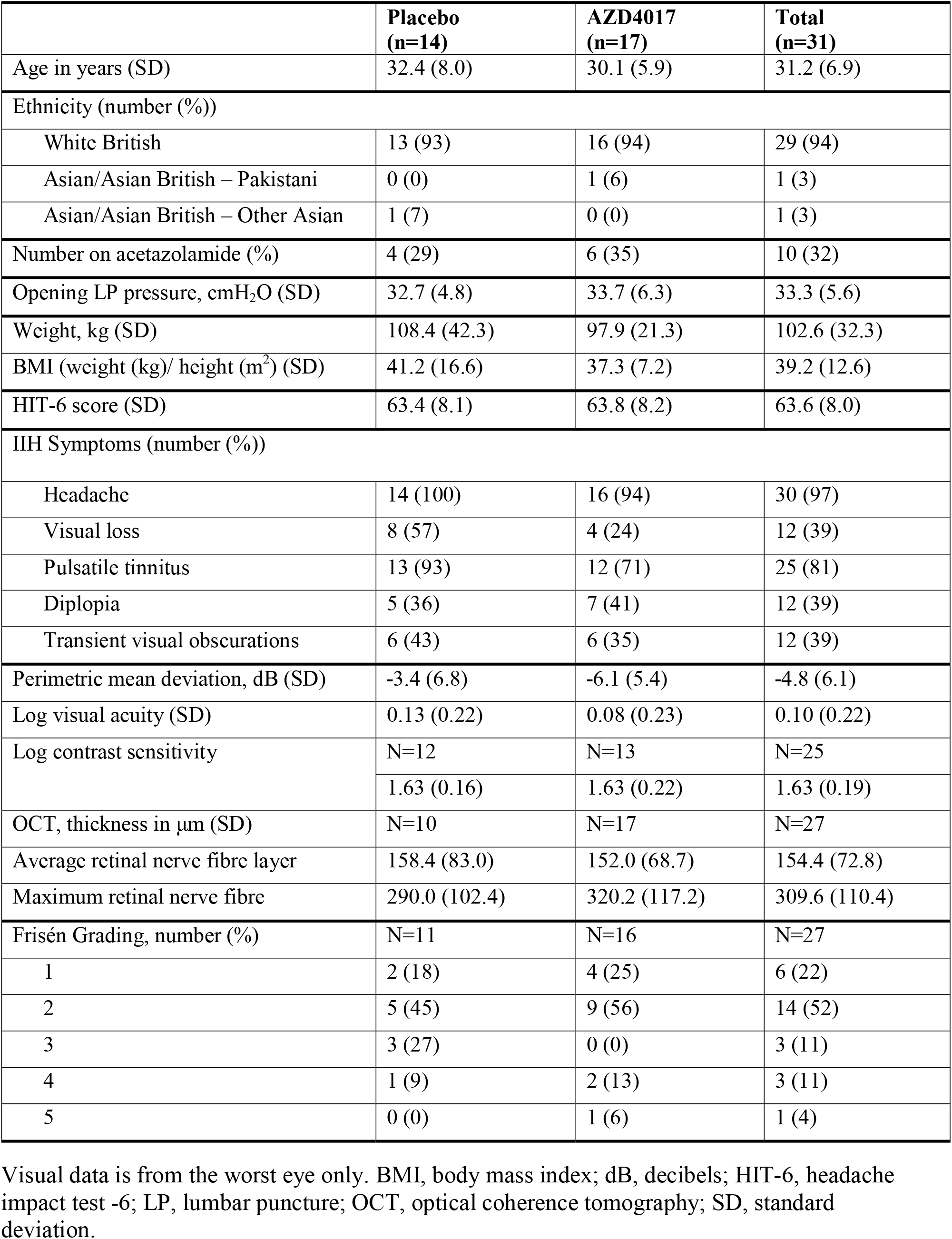
Baseline characteristics and ophthalmic measurements

|  | **Placebo**<br>**(n=14)** | **AZD4017**<br>**(n=17)** | **Total**<br>**(n=31)** |
| --- | --- | --- | --- |
| Age in years (SD) | 32.4 (8.0) | 30.1 (5.9) | 31.2 (6.9) |
| Ethnicity (number (%)) |  |  |  |
| White British | 13 (93) | 16 (94) | 29 (94) |
| Asian/Asian British – Pakistani | 0 (0) | 1 (6) | 1 (3) |
| Asian/Asian British – Other Asian | 1 (7) | 0 (0) | 1 (3) |
| Number on acetazolamide (%) | 4 (29) | 6 (35) | 10 (32) |
| Opening LP pressure, cmH<sub>2</sub>O (SD) | 32.7 (4.8) | 33.7 (6.3) | 33.3 (5.6) |
| Weight, kg (SD) | 108.4 (42.3) | 97.9 (21.3) | 102.6 (32.3) |
| BMI (weight (kg)/ height (m<sup>2</sup>) (SD) | 41.2 (16.6) | 37.3 (7.2) | 39.2 (12.6) |
| HIT-6 score (SD) | 63.4 (8.1) | 63.8 (8.2) | 63.6 (8.0) |
| IIH Symptoms (number (%)) |  |  |  |
| Headache | 14 (100) | 16 (94) | 30 (97) |
| Visual loss | 8 (57) | 4 (24) | 12 (39) |
| Pulsatile tinnitus | 13 (93) | 12 (71) | 25 (81) |
| Diplopia | 5 (36) | 7 (41) | 12 (39) |
| Transient visual obscurations | 6 (43) | 6 (35) | 12 (39) |
| Perimetric mean deviation, dB (SD) | -3.4 (6.8) | -6.1 (5.4) | -4.8 (6.1) |
| Log visual acuity (SD) | 0.13 (0.22) | 0.08 (0.23) | 0.10 (0.22) |
| Log contrast sensitivity | N=12 | N=13 | N=25 |
|  | 1.63 (0.16) | 1.63 (0.22) | 1.63 (0.19) |
| OCT, thickness in μm (SD) | N=10 | N=17 | N=27 |
| Average retinal nerve fibre layer | 158.4 (83.0) | 152.0 (68.7) | 154.4 (72.8) |
| Maximum retinal nerve fibre | 290.0 (102.4) | 320.2 (117.2) | 309.6 (110.4) |
| Frisén Grading, number (%) |  |  |  |
| 1 | 2 (18) | 4 (25) | 6 (22) |
| 2 | 5 (45) | 9 (56) | 14 (52) |
| 3 | 3 (27) | 0 (0) | 3 (11) |
| 4 | 1 (9) | 2 (13) | 3 (11) |
| 5 | 0 (0) | 1 (6) | 1 (4) |

Visual data is from the worst eye only. BMI, body mass index; dB, decibels; HIT-6, headache impact test -6; LP, lumbar puncture; OCT, optical coherence tomography; SD, standard deviation.

### Clinical Outcomes

#### Primary Clinical Outcome

At 12 weeks, the mean ICP was 29.7 cmH_2_O (SD=5.2) in the AZD4017 group compared with 31.3 cmH_2_O (SD=6.7) in the placebo group (adjusted mean difference:-2.8 cmH_2_O, 95% confidence interval (CI): −7.1-1.5; p=0.2) (Figure 2A). An exploratory analysis assessed the mean change in ICP within each group. ICP decreased from 33.7 (SD=6.3) at baseline to 29.7 cmH_2_O (SD=5.2) at 12 weeks in the AZD4017 group (mean change: −4.3 cmH_2_O (SD=5.7); *p*=0.009) and from 32.7 (SD=4.8) to 31.3 cmH_2_O (SD=6.7) in the placebo group (mean change: −0.3 cmH_2_O (SD=5.9); p=0.8 (Figure 2B,C).

**Figure 2.**
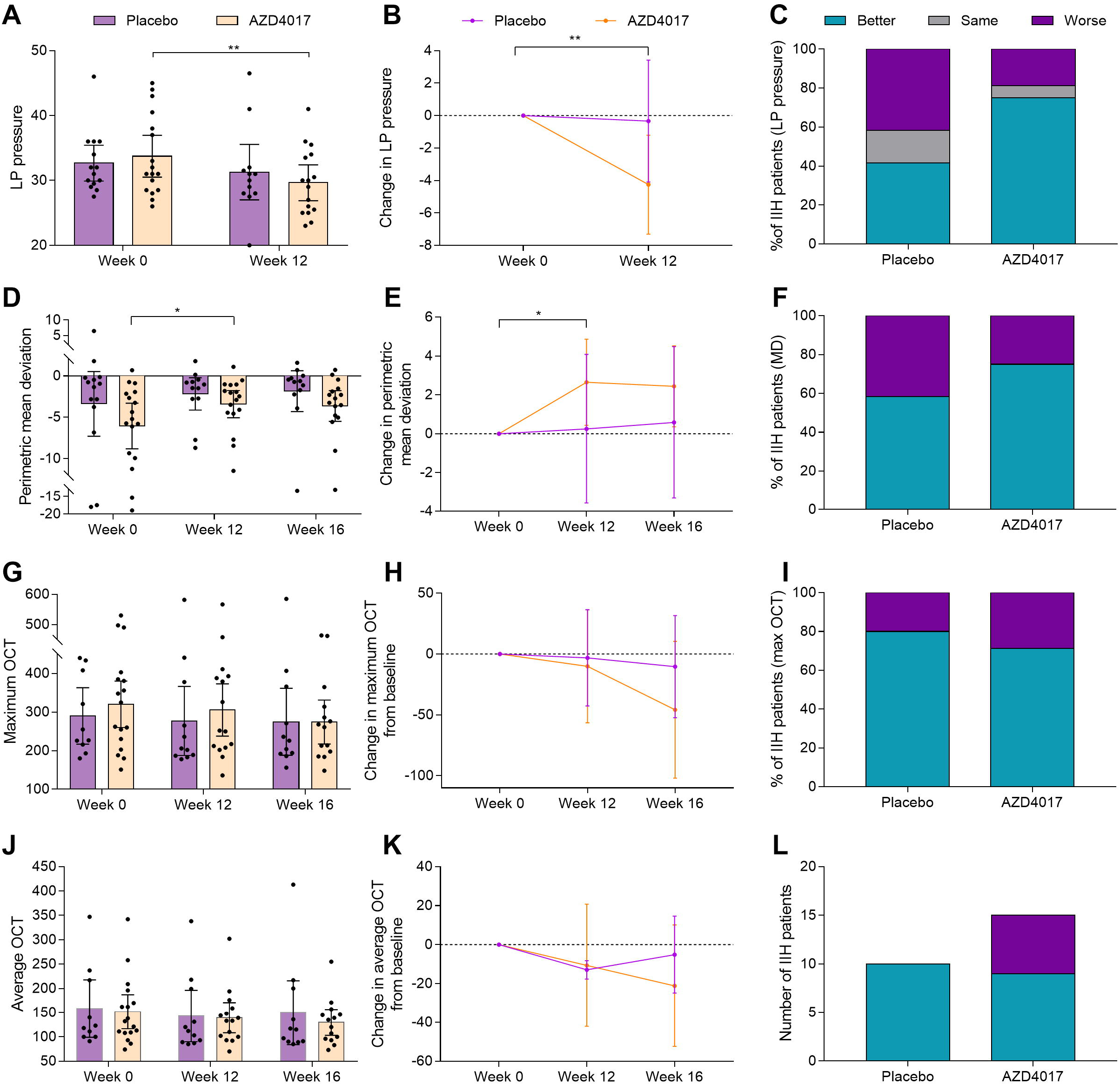
Clinical outcomes following treatment with AZD4017 and placebo for 12 weeks and then 4 weeks after stopping treatment. **(A)** Absolute LP pressure. **(B)** Change in LP pressure. **(C)** Percentage of patients with better, same or worse LP pressure at 12 weeks. **(D)** Absolute visual field mean deviation (dB). **(E)** Change in visual field mean deviation. **(F)** Percentage of patients with better, same or worse visual field mean deviation at 12 weeks. **(G)** Absolute maximum optical coherence tomography (OCT) retinal nerve fibre layer (RNFL) height (µm). **(H)** Change in maximum OCT RNFL height. **(I)** Percentage of patients with better, same or worse maximum OCT RNFL height at 12 weeks. **(J)** Average OCT RNFL height (µm). **(K)** Change in average OCT RNFL height. **(L)** Number of patients with better, same or worse average OCT RNFL height at 12 weeks. Data is presented as mean ± 95% confidence index. *<0.05, ** < 0.01.

#### Secondary Clinical Outcomes

At weeks 12 and 16, there were no statistically significant differences between the two treatment groups in IIH symptoms (Suppl. Table 2). At 12 and 16 weeks, the Humphrey Visual Field Analyzer PMD (worst eye) was not significantly different between groups (adjusted mean difference at 12 weeks: 0.3dB, 95% CI: −2.0-2.7, *p*=0.8) (Figure 2D, E, F; Table 2, Suppl. Table 3). However, within group analysis showed that the PMD improved from −6.1 dB (SD=5.4) at baseline to −3.4 dB (SD=3.2) (mean change 2.7 dB (SD=4.3), p=0.04) at 12 weeks in the AZD4017 group and from −3.4 dB (SD=6.8) to −2.2 dB (SD=3.1) (mean change 0.3 dB (SD=6.0), p=1.0) in the placebo group. There were also no statistically significant differences between groups at either 12 or 16 weeks in visual acuity, contrast sensitivity, OCT average and maximal RNFL (Table 2; Figure 2G, H,I maximum RNFL, Figure 2J,K,L average RNFL, Suppl. Table 3). At 12 weeks, the mean Frisén grade in the worst eye was 1.56 (SD=0.96) in the AZD4017 group and 2.25 (SD=0.87) in the placebo group (adjusted mean difference: −0.7, 95% CI:-1.4-0.3; p=0.06).

**Table 2:** Visual function and optic nerve head at baseline and week 12

|  | **Baseline**<br>**Mean value (SD)** |  | **Week 12**<br>**Mean value (SD)** |  | **Adjusted**<br>**mean**<br>**difference at**<br>**12 weeks**<br>**(95% C.I)** | **P value** |
| --- | --- | --- | --- | --- | --- | --- |
|  | **Placebo** | **AZD4017** | **Placebo** | **AZD4017** |  |  |
| **Worse Eye** |  |  |  |  |  |  |
| **Visual Acuity**<br>**LogMAR** | 0.13<br>(0.22) | 0.08<br>(0.23) | 0.09<br>(0.18) | 0.06<br>(0.15) | -0.03<br>(-0.12, 0.07) | 0.5 |
| **Contrast**<br>**Sensitivity** | 1.63<br>(0.16) | 1.63<br>(0.22) | 1.66<br>(0.12) | 1.65<br>(0.15) | -0.02<br>(-0.15, 0.11) | 0.7 |
| **Perimetric**<br>**Mean Deviation** | -3.4<br>(6.8) | -6.1<br>(5.4) | -2.2<br>(3.1) | -3.4<br>(3.2) | 0.3<br>(-2.0, 2.7) | 0.8 |
| **OCT RNFL**<br>**Average (µm)** | 158.4<br>(83.0) | 152.0<br>(68.7) | 143.2<br>(78.7) | 139.7<br>(56.3) | 0.1<br>(-34.0, 34.1) | 1.0 |
| **OCT Maximal**<br>**RNFL (µm)** | 290.0<br>(102.4) | 320.2<br>(117.2) | 277.0<br>(133.1) | 305.5<br>(122.3) | -4.5<br>(-68.1, 59.1) | 0.9 |
| **Average Frisén**<br>**grading** | 2.27<br>(0.90) | 2.19<br>(1.17) | 2.25<br>(0.87) | 1.56<br>(0.96) | -0.7<br>(-1.4, 0.03) | 0.06 |

All measures shown in the table are of worst eye. Negative values in the adjusted mean difference between treatment arms favour AZD4017. C.I: confidence interval; OCT: optical coherence tomography; RNFL: retinal nerve fibre layer; SD: standard deviation.

All headache outcomes were not statistically significantly different between AZD4017 and placebo at weeks 12 or 16 (Suppl. Table 4). There were also no statistically significant differences in any of the anthropological outcomes (BMI, waist:hip ratio).

### Safety and Tolerability

Study medication was well tolerated with participants in both arms taking on average 98% of the total 168 study medication doses (mean doses taken were 164 (range 146 – 168) and 165 (range 158-168) in the AZD4017 and placebo group respectively). There were no participant withdrawals due to adverse effects. Nine adverse events (in 6 participants) were deemed related to AZD4017, none were serious and 3 were due to non-clinically relevant fluctuations in serum cortisol. Adverse events are shown in Suppl. Table 5. One serious adverse event was reported in the placebo arm and deemed unrelated (fulminant deterioration in IIH necessitating CSF shunting one day post-randomization).

No differences were noted between treatment groups for the safety blood tests (Suppl. Table 6). As expected, there was a rise in the HPA stimulatory hormone, ACTH, over 12 weeks in the AZD4017 group (mean difference at 12 weeks: 12.36ng/l, 95%CI: −0.03-24.74). There was no difference in serum cortisol, testosterone or androstenedione, although serum DHEAS, a marker of adrenal androgen production was higher at 12 weeks in the AZD4017 group (mean difference at 12 weeks: 5.44nmol/l, 95%CI, 1.09-9.79); levels returned to normal 4 weeks after treatment cessation (week 16).

### In vivo assessments

#### Blood and CSF levels of AZD4017 and glucocorticoids

AZD4017 concentrations were detected in the serum after one week of treatment and sustained at week 12 (n=6). The presence of AZD4017 in the CSF was 0.5% that of the serum (Suppl. Table 7). No AZD4017 was detected in the placebo group at any time point. There was no correlation between serum or CSF drug levels and LP pressure (serum: *r*=0.03; *p*=1.0; CSF: *r*=−0.2; *p*=0.7) or visual field mean deviation (serum: *r*=-0.2; *p*=0.7; CSF: *r*=-0.04; *p*=0.9). Serum and CSF cortisol and cortisone were examined in the placebo and AZD4017 groups at baseline and at 12 weeks. The serum cortisol:cortisone ratio was not significantly different between arms at weeks 0 (*p*=0.6) and 12 (*p*=0.5) (Figure 3A). The CSF cortisol:cortisone ratio did not differ between arms at baseline (*p*=0.9); however, at week 12 there was a significant decrease in the CSF cortisol:cortisone in the AZD4017 group compared to placebo (*p*=0.002), and the AZD4017 group between baseline and 12 weeks (*p*=0.03) (Figure 3B), implying local 11β-HSD1 activity can regulate CSF glucocorticoid exposure.

**Figure 3.**
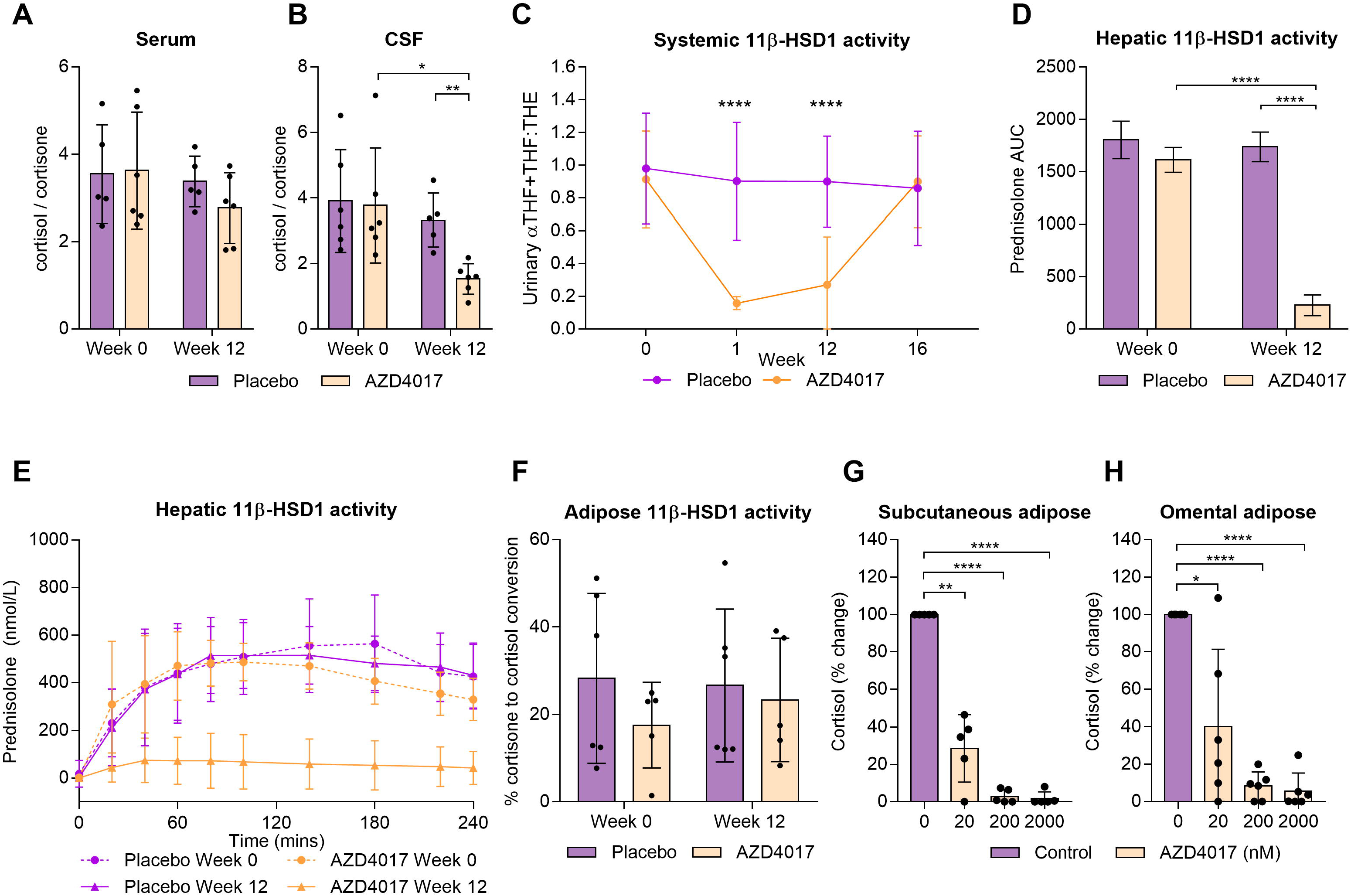
*In vivo* and *ex vivo* analysis of 11β-HSD activity after 12 weeks treatment with either AZD4017 or placebo. **(A)** Serum cortisol:cortisone ratio. **(B)** CSF cortisol:cortisone ratio. **(C)** Urinary 11β-HSD1 activity ((5α-THF+THF):THE) at weeks 0, 1, 12 and 16. **(D)** Change in prednisolone area under the curve (AUC) (see E). **(E)** Hepatic 11β-HSD1 activity (mean blood prednisolone concentration after conversion from prednisone) over 4 hours. **(F)** Subcutaneous adipose 11β-HSD1 activity (percentage change from cortisone to cortisol) *ex vivo.* **(G)** *Ex vivo* subcutaneous adipose **(H)** Omental adipose 11β-HSD1 activity (cortisol production from cortisone) after 24 hours incubation with either 0, 20, 200 or 2000 nM of AZD4017 *in vitro.* Data presented as mean±SD. * p<0.05, ** p<0.01, ***p<0.001, ****p<0.0001

#### *In vivo* systemic 11**β**-Hydroxysteroid Dehydrogenase activity

The urinary (5αTHF+THF):THE glucocorticoid metabolite ratio reflective of systemic 11β-HSD1 activity was significantly reduced in AZD4017 vs. placebo groups at week 1 (0.16±0.04 versus 0.90±0.36, *p*<0.0001) and week 12 (0.27±0.29 versus 0.90±0.28; *p*<0.0001). By contrast, the ratios did not differ between the two treatment groups at baseline (*p*=0.6) and 4 weeks after the end of treatment (week 16, *p*=0.8). 11β-HSD type 2 activity as assessed by urinary cortisol over cortisone remained unchanged and similar in both groups throughout the 12 weeks of treatment (*p*=0.6). These data imply that AZD4017 was effective at inhibiting 11β-HSD1 (Figure 3C). No correlation was found between the change in (5αTHF+THF):THE and ICP (*r*=0.1; *p*=0.7) or PMD (*r*=0.2; *p*=0.4).

#### Hepatic 11**β**-Hydroxysteroid Dehydrogenase activity

The placebo group had robust capacity to generate prednisolone following oral prednisone at both baseline and after 12 weeks. The baseline prednisolone generation curve for the AZD4017 group was indistinguishable from the placebo curve; however at 12 weeks the AZD4017 group were essentially unable to generate prednisolone (Figure 3D,E), indicating effective inhibition of hepatic 11β-HSD1 activity. Area under the curve analysis of the mean time points at 12 weeks showed significantly impaired prednisolone generating capacity for AZD4017 vs. placebo (228±99 vs 1738±142; *p*<0.0001), an 85.9% reduction (*p*<0.0001) in overall prednisolone generating capacity after 12 weeks (Figure 3D,E). There was no correlation between the change in the area under the curve for prednisolone and ICP (*r*=0.1; *p*=0.8) or PMD (*r*=0.4; *p*=0.2).

#### Adipose 11**β**-Hydroxysteroid Dehydrogenase activity

While AZD4017 effectively inhibited hepatic 11β-HSD1, we were unable to show impaired capacity to generate cortisol from cortisone in explanted subcutaneous adipose biopsies. At baseline and following 12 weeks of oral AZD4017, there was no significant change in total cortisol vs. placebo (9.0±5.6 vs 12.4±4.9 nmol; *p*=0.3) or percentage conversion of cortisone to cortisol (23±14 vs 27±18 %; *p*>0.99) (Figure 3F). However, AZD4017 was able to significantly inhibit 11β-HSD1 activity when added to *ex vivo* adipose explants from subcutaneous and omental depots. 20nM AZD4017 significantly impaired conversion of cortisone to cortisol (>70% vs. control), 200nM onwards was sufficient to effectively block cortisol generation, particularly in the subcutaneous depot (Figure 3G,H).

## Discussion

We report the first phase 2 RCT assessing 11β-HSD1 inhibitor AZD4017 for the treatment of IIH. We have shown some possible clinical benefit for AZD4017, and that it was well tolerated and safe. We found evidence for effective *in vivo* 11β-HSD1 inhibition.

Our primary hypothesis stated that 11β-HSD1 inhibition in IIH patients would reduce CSF secretion and lower ICP while being safe and tolerable following 12 weeks of treatment. ICP was the primary clinical outcome measure, representing the hallmark of the disease driving clinical sequelae. At 12 weeks, although ICP was lower in the AZD4017 group compared to placebo, the difference between groups was not statistically significant. Exploratory analyses of the mean change within groups found a significant improvement in ICP in the AZD4017 group between baseline and 12 weeks, but not in the placebo group. Previous trials have noted that ICP reduction below the cut off of 25 cmH_2_O is not universally required to translate into resolution of IIH clinical features.^(Sinclair *et al.*, 2010b)^ A minimal clinically important change in ICP in IIH has not been determined, and establishing one would be useful for future trials.

The visual field perimetric assessment is another clinically meaningful measure and has been selected as the primary outcome measures in previous IIH trials. We found no difference between groups in PMD at 12 weeks, however, there was significant improvement over time in the AZD4017 arm but not in the placebo arm. This may reflect the pragmatic recruitment of all degrees of PMD at enrolment whilst other trials have enrolled a selected cohort (e.g. −2 to −5 dB).(Wall et al., 2014) Additionally, his small trial was not powered to determine significant in the secondary outcome measures.

Headache is a key disabling feature in IIH.(Mulla *et al.*, 2015) We did not detect differences between the groups in any of the headache assessments at 12 weeks, although data from the patient completed HIT-6 favoured the AZD4017 group. Evaluating the effect of AZD4017 on headache measures over a longer treatment duration would be of interest.

Previous trials showed that 11β-HSD1 inhibition leads to adaptive changes in HPA stimulatory hormone ACTH and the adrenal androgen precursor DHEA. Our data support these findings, but with no significant change in downstream effector hormones (cortisol and testosterone).

*In vivo* evaluation of our patients demonstrated that AZD4017 was a highly effective systemic and hepatic 11β-HSD1 inhibitor, in line with previous studies using11β-HSD1 inhibitors in humans(Schwab *et al.*, 2017)’(Courtney *et al.*, 2008). Systemic efficacy may modify metabolic aspects of IIH with indirect benefits on ICP.(Hornby *et al.*, 2018)

While AZD4017 effectively inhibited 11β-HSD1 when applied to subcutaneous and omental adipose tissue explants, we were unable to prove inhibition *in vivo*, and propose that 11β-HSD1 activity recovers over the assay period once removed from AZD4017, a reversible competitive inhibitor.

Blood-brain-barrier AZD4017 penetrance was low, with levels in the CSF 0.5% those of plasma levels, but were associated with reduced CSF cortisol:cortisone ratio suggesting that 11β-HSD1 may contribute to cortisol availability in the CSF. The unchanged serum cortisol:cortisone is likely reflective of HPA axis set-point.

## Limitations

We were unable to directly evaluate 11β-HSD1 inhibition at the choroid plexus, the tissue responsible for CSF secretion. We have evaluated efficacy of other IIH drugs using rodent ICP monitoring models,(Botfield *et al.*, 2017; Scotton *et al.*, 2018) but AZD4017 is only effective in humans and primates, thus limiting our ability to evaluate its action in rodent models. The trial duration was likely too short. A duration of 12 weeks was chosen for evaluation of safety and tolerability and represented the longest duration of dosing to date with AZ4017. This may not have been sufficient for meaningful evaluation of clinical outcomes with other IIH RCTs evaluating drugs over a 6 month period.(Committee *et al.*, 2014) The enrolment criteria for the study were deliberately broad allowing inclusion of a spectrum of IIH patients with active disease and ensuring generalizability of results; however, this did not allow evaluation in disease subgroups such as those with mild visual loss vs. those with severe irreversible visual loss. Finally, the sample size (31 participants) is small which may have limited meaningful evaluation of clinical measures and the trial was not design to establish significant changes in the secondary clinicaloutcome measures.

## Conclusions

This is the first phase II study evaluating the novel pharmacological therapy AZD4017 in IIH. We demonstrate safety, tolerability and provide strong *in vivo* evidence for effective 11β-HSD1 inhibition. There was a significant reduction in ICP in the AZD4017 and not the placebo group over the treatment duration (exploratory within group analysis); however, the primary analysis evaluating the difference between groups at 12 weeks did not reach statistical significance. The data suggest that 11β-HSD1 inhibition may have utility for reducing the effects and consequences of raised ICP in patients with IIH. Further evaluation of these therapeutic strategies in this disabling disease, for which few useful medical options exist, would be worthwhile.

## Abbreviations

11β-: 11β-hydroxysteroid dehydrogenase type 1
HSD1: inhibitor
5αTHF: 5α-tetrahydrocortisol
ACTH: Adrenocorticotropic hormone
BBB: Blood brai barrier
CP: Choroid Plexus
CSF: Cerebrospinal fluid
DHEAS: Dehydroepiandrosterone sulfate
E: Cortisone
F: Cortisol
HPA: Hypothalamic pituitary adrenal
ICP: intracranial pressure
IIH: Idiopathic Intracranial Hypertension
LC-: liquid chromatography-tandem mass
MS/MS: spectrometry
OCT: Optical coherence tomography
PMD: Perimetric mean deviation
RCT: Randomised Controlled Trial
RNFL: Retinal nerve fibre layer
THE: tetrahydrocortisone
THF: tetrahydrocortisol
VA: Visual acuity

## Contributions

KM, JM & HB made major contributions to the acquisition, and interpretation of data; and drafting of the work.

RW & NI made substantial contributions to the clinical statistics and undertook the data analysis of the clinical data.

SM, RO, TM, AK, PS, CR, WS & AY made substantial contributions to the clinical data acquisition and drafting of the work.

AT, LG, CJ & WA were responsible for steroid analysis in serum, CSF and tissue by LC-MS/MS analysis and contributed to the drafting of the work.

RS & CW made substantial contributions to the adipose data acquisition and drafting of the work.

RF made a substantial contribution to study safety and delivery.

PS, JT & GL were involved in conceptualization of the hypothesis and drafting and revision of the work.

AS conceptualized the hypothesis and led on study design, oversight, interpretation of data and drafting of the work.

All authors approved the final version to be submitted for review.

## Acknowledgements

We acknowledge Birmingham Clinical Trials Unit for trial coordination, data management and clinical data analysis. We thank Peter Nightingale, Statistician, NIHR/Wellcome Trust Clinical Research Facility, University Hospitals Birmingham NHS Foundation Trust, Birmingham, B15 2TH, UK, for help with the exploratory analyses. We acknowledge the support of the National Institute of Health Research Clinical Research Network (NIHR CRN) and the nurses and staff of the NIHR/Wellcome Trust Clinical Research Facilities where IIH:DT was performed. The views expressed in this publication are those of the authors and not necessarily those of the MRC, NIHR or the Department of Health. We thank AstraZeneca for the AZD4017 compound and for their helpful advice, specifically from Madeleine Brady, K. Jane Escott, Rebecca J Fairclough, Alison Holt, James Sylvester, Lorraine C. Webber and Chris Wilks. We acknowledge Almac Group, UK for the randomization.

## Supplemental files

Supplemental File 1: Additional data

## References

AstraZeneca. Phase I Study in Healthy Volunteers to Assess the Safety, Tolerability, Pharmacokinetics and Pharmacodynamics of AZD4017 After Repeated Ascending Oral Doses (MAD). 2000-[01 March 2017]-a [cited; Available from: http://clinicaltrials.gov/show/NCT00841048

AstraZeneca. A phase IIa study to assess the tolerability, safety and efficacy of AZD4017 for raised intra-ocular pressure. 2000-[01 March 2017]-b [cited; Available from: http://clinicaltrials.gov/show/NCT01173471

AstraZeneca. Study to Investigate Safety and Tolerability Single Ascending Doses of AZD4017. 2000-[01 March 2017]-c [cited; Available from: http://clinicaltrials.gov/show/NCT00791752

AstraZeneca. Study to Investigate Safety and Tolerability Single Ascending Doses of AZD4017. 2000-[01 March 2017]-d [cited; Available from: http://clinicaltrials.gov/show/NCT00799747

AstraZeneca. Study to Evaluate Methods That Assess the Effect of AZD4017 in Adipose Tissue. 2000-[01 March 2017]. [cited; Available from: http://clinicaltrials.gov/show/NCT01096004

Ball AK, Howman A, Wheatley K, Burdon MA, Matthews T, Jacks AS, et al. A randomised controlled trial of treatment for idiopathic intracranial hypertension. J Neurol 2011; 258(5): 874–81.

Bayliss MS, Bjornerm JB, Kosinski M, Dahlöf CGH, Dowson A, Cady RK. Development of HIT-6, a paper-based short form for measuring headache impact. In: Olesen J, Steiner TJ, Lipton RB Frontiers in Headache Research Reducing the Burden of Headache New York: Oxford University Press-USA 2003; 11: 386–90.

Botfield HF, Uldall MS, Westgate CSJ, Mitchell JL, Hagen SM, Gonzalez AM, et al. A glucagon-like peptide-1 receptor agonist reduces intracranial pressure in a rat model of hydrocephalus. Sci Transl Med 2017; 9(404).

Boyle CD. Recent advances in the discovery of 11beta-HSD1 inhibitors. Curr Opin Drug Discov Devel 2008; 11(4): 495–511.

Colquitt JL, Pickett K, Loveman E, Frampton GK. Surgery for weight loss in adults. Cochrane Database of Systematic Reviews 2014(8).

Committee NIIHSGW, Wall M, McDermott MP, Kieburtz KD, Corbett JJ, Feldon SE, et al. Effect of acetazolamide on visual function in patients with idiopathic intracranial hypertension and mild visual loss: the idiopathic intracranial hypertension treatment trial. JAMA 2014; 311(16): 1641–51.

Courtney R, Stewart PM, Toh M, Ndongo M-N, Calle RA, Hirshberg B. Modulation of 11β-Hydroxysteroid Dehydrogenase (11βHSD) Activity Biomarkers and Pharmacokinetics of PF-00915275, a Selective 11βHSD1 Inhibitor. The Journal of Clinical Endocrinology & Metabolism 2008; 93(2): 550–6.

Daniels AB, Liu GT, Volpe NJ, Galetta SL, Moster ML, Newman NJ, et al. Profiles of obesity, weight gain, and quality of life in idiopathic intracranial hypertension (pseudotumor cerebri). Am J Ophthalmol 2007; 143(4): 635–41.

Davson H. Formation and drainage of the cerebrospinal fluid. The Scientific basis of medicine annual reviews 1966: 238–59.

Eftekhari S, Salvatore CA, Johansson S, Chen T-b, Zeng Z, Edvinsson L. Localization of CGRP, CGRP receptor, PACAP and glutamate in trigeminal ganglion. Relation to the blood–brain barrier. Brain Research 2015; 1600: 93–109.

Friedman DI, Liu GT, Digre KB. Revised diagnostic criteria for the pseudotumor cerebri syndrome in adults and children. Neurology 2013; 81(13): 1159–65.

Friedman DI, McDermott MP, Kieburtz K, Kupersmith M, Stoutenburg A, Keltner JL, et al. The idiopathic intracranial hypertension treatment trial: design considerations and methods. J Neuroophthalmol 2014; 34(2): 107–17.

Frisen L. Swelling of the optic nerve head: a staging scheme. J Neurol Neurosurg Psychiatry 1982; 45(1): 13–8.

Gathercole LL, Lavery GG, Morgan SA, Cooper MS, Sinclair AJ, Tomlinson JW, et al. 11β-Hydroxysteroid Dehydrogenase 1: Translational and Therapeutic Aspects. Endocrine Reviews 2013; 34(4): 525–55.

Hassan-Smith ZK, Morgan SA, Sherlock M, Hughes B, Taylor AE, Lavery GG, et al. Gender-Specific Differences in Skeletal Muscle 11beta-HSD1 Expression Across Healthy Aging. The Journal of clinical endocrinology and metabolism 2015; 100(7): 2673–81.

Hoffmann J, Mollan SP, Paemeleire K, Lampl C, Jensen RH, Sinclair AJ. European headache federation guideline on idiopathic intracranial hypertension. The journal of headache and pain 2018; 19(1): 93.

Hornby C, Mollan SP, Botfield H, O’Reilly MW, Sinclair AJ. Metabolic Concepts in Idiopathic Intracranial Hypertension and Their Potential for Therapeutic Intervention. J Neuroophthalmol 2018.

Juhlen R, Idkowiak J, Taylor AE, Kind B, Arlt W, Huebner A, et al. Role of ALADIN in human adrenocortical cells for oxidative stress response and steroidogenesis. PloS one 2015; 10(4): e0124582.

Long JE, Drayson MT, Taylor AE, Toellner KM, Lord JM, Phillips AC. Morning vaccination enhances antibody response over afternoon vaccination: A cluster-randomised trial. Vaccine 2016; 34(24): 2679–85.

Manfield JH, Yu KKH, Efthimiou E, Darzi A, Athanasiou T, Ashrafian H. Bariatric Surgery or Non-surgical Weight Loss for Idiopathic Intracranial Hypertension? A Systematic Review and Comparison of Meta-analyses. Obesity Surgery 2017; 27(2): 513–21.

Markey KA, Mollan SP, Jensen RH, Sinclair AJ. Understanding idiopathic intracranial hypertension: mechanisms, management, and future directions. The Lancet Neurology 2016; 15(1): 78–91.

Markey KA, Ottridge R, Mitchell JL, Rick C, Woolley R, Ives N, et al. Assessing the Efficacy and Safety of an 11beta-Hydroxysteroid Dehydrogenase Type 1 Inhibitor (AZD4017) in the Idiopathic Intracranial Hypertension Drug Trial, IIH:DT: Clinical Methods and Design for a Phase II Randomized Controlled Trial. JMIR research protocols 2017a; 6(9): e181.

Markey KA, Ottridge R, Mitchell JL, Rick C, Woolley R, Ives N, et al. Assessing the Efficacy and Safety of an 11β-Hydroxysteroid Dehydrogenase Type 1 Inhibitor (AZD4017) in the Idiopathic Intracranial Hypertension Drug Trial, IIH:DT: Clinical Methods and Design for a Phase II Randomized Controlled Trial. JMIR Research Protocols 2017b; 6(9): e181.

Mollan SP, Aguiar M, Evison F, Frew E, Sinclair AJ. The expanding burden of idiopathic intracranial hypertension. Eye (London, England) 2018a.

Mollan SP, Davies B, Silver NC, Shaw S, Mallucci CL, Wakerley BR, et al. Idiopathic intracranial hypertension: consensus guidelines on management. J Neurol Neurosurg Psychiatry 2018b.

Mollan SP, Hornby C, Mitchell J, Sinclair AJ. Evaluation and management of adult idiopathic intracranial hypertension. Practical neurology 2018c.

Mollan SP, Markey KA, Benzimra JD, Jacks A, Matthews TD, Burdon MA, et al. A practical approach to, diagnosis, assessment and management of idiopathic intracranial hypertension. Pract Neurol 2014.

Mooij CF, Parajes S, Rose IT, Taylor AE, Bayraktaroglu T, Wass JA, et al. Characterization of the molecular genetic pathology in patients with 11beta-hydroxylase deficiency. Clinical endocrinology 2015; 83(5): 629–35.

Mulla Y, Markey KA, Woolley RL, Patel S, Mollan SP, Sinclair AJ. Headache determines quality of life in idiopathic intracranial hypertension. The journal of headache and pain 2015; 16: 521.

O’Reilly MW, Westgate CS, Hornby C, Botfield H, Taylor AE, Markey K, et al. A unique androgen excess signature in idiopathic intracranial hypertension is linked to cerebrospinal fluid dynamics. JCI insight 2019.

Piper RJ, Kalyvas AV, Young AM, Hughes MA, Jamjoom AA, Fouyas IP. Interventions for idiopathic intracranial hypertension. The Cochrane database of systematic reviews 2015(8): Cd003434.

Rauz S, Cheung CM, Wood PJ, Coca-Prados M, Walker EA, Murray PI, et al. Inhibition of 11beta-hydroxysteroid dehydrogenase type 1 lowers intraocular pressure in patients with ocular hypertension. Qjm 2003; 96(7): 481–90.

Richards J, Lim AC, Hay CW, Taylor AE, Wingate A, Nowakowska K, et al. Interactions of abiraterone, eplerenone, and prednisolone with wild-type and mutant androgen receptor: a rationale for increasing abiraterone exposure or combining with MDV3100. Cancer research 2012; 72(9): 2176–82.

Sagmeister MS, Taylor AE, Fenton A, Wall NA, Chanouzas D, Nightingale PG, et al. Glucocorticoid activation by 11beta-hydroxysteroid dehydrogenase enzymes in relation to inflammation and glycaemic control in chronic kidney disease: a cross-sectional study. Clinical endocrinology 2018.

Sandeep TC, Andrew R, Homer NZ, Andrews RC, Smith K, Walker BR. Increased in vivo regeneration of cortisol in adipose tissue in human obesity and effects of the 11beta-hydroxysteroid dehydrogenase type 1 inhibitor carbenoxolone. Diabetes 2005; 54(3): 872–9.

Schwab D, Sturm C, Portron A, Fuerst-Recktenwald S, Hainzl D, Jordan P, et al. Oral administration of the 11β-hydroxysteroid-dehydrogenase type 1 inhibitor RO5093151 to patients with glaucoma: an adaptive, randomised, placebo-controlled clinical study. BMJ Open Ophthalmology 2017; 1(1).

Scotton WJ, Botfield HF, Westgate CS, Mitchell JL, Yiangou A, Uldall MS, et al. Topiramate is more effective than acetazolamide at lowering intracranial pressure. Cephalalgia: an international journal of headache 2018: 333102418776455.

Sinclair A, Burdon M, Ball A, Nightingale N, Good P, Matthews T, et al. Low energy diet and intracranial pressure in women with idiopathic intracranial hypertension: prospective cohort study. BMJ 2010a; 7: 341.

Sinclair AJ, Burdon MA, Nightingale PG, Ball AK, Good P, Matthews TD, et al. Low energy diet and intracranial pressure in women with idiopathic intracranial hypertension: prospective cohort study. BMJ 2010b; 341: c2701.

Sinclair AJ, Onyimba CU, Khosla P, Vijapurapu N, Tomlinson JW, Burdon MA, et al. Corticosteroids, 11beta-Hydroxysteroid Dehydrogenase Isozymes and the Rabbit Choroid Plexus. J Neuroendocrinol 2007; 19(8): 614–20.

Sinclair AJ, Walker EA, Burdon MA, van Beek AP, Kema IP, Hughes BA, et al. Cerebrospinal fluid corticosteroid levels and cortisol metabolism in patients with idiopathic intracranial hypertension: a link between 11beta-HSD1 and intracranial pressure regulation? The Journal of clinical endocrinology and metabolism 2010c; 95(12): 5348–56.

Stefan N, Ramsauer M, Jordan P, Nowotny B, Kantartzis K, Machann J, et al. Inhibition of 11beta-HSD1 with RO5093151 for non-alcoholic fatty liver disease: a multicentre, randomised, double-blind, placebo-controlled trial. The lancet Diabetes & endocrinology 2014; 2(5): 406–16.

Tomlinson JW, Stewart PM. Cortisol metabolism and the role of 11beta-hydroxysteroid dehydrogenase. Best Pract Res Clin Endocrinol Metab 2001; 15(1): 61–78.

Wake DJ, Walker BR. 11 beta-hydroxysteroid dehydrogenase type 1 in obesity and the metabolic syndrome. Mol Cell Endocrinol 2004; 215(1-2): 45–54.

Wall M, McDermott MP, Kieburtz KD, Corbett JJ, Feldon SE, Friedman DI, et al. Effect of acetazolamide on visual function in patients with idiopathic intracranial hypertension and mild visual loss: the idiopathic intracranial hypertension treatment trial. Jama 2014; 311(16): 1641–51.

