## Supplemental data for "11β-Hydroxysteroid Dehydrogenase Type 1 inhibition in Idiopathic Intracranial Hypertension: a double-blind randomized controlled trial"

**Table 1 supplemental: Eligibility criteria**

| Inclusion Criteria | |
| --- | --- |
| 1. | Provision of informed consent |
| 2. | Female patients between 18 and 55 years |
| 3. | Diagnosis of IIH by the Modified Dandy criteria^18^ with active disease (papilledema (Frisen grade greater than or equal to 1) and significantly raised ICP > 25cmH_2_O) and normal brain imaging (including magnetic resonance venography or CT with venography). |
| 4. | Participants must be willing to use one form of highly effective non-hormonal contraception. |
| 5. | Participants are able to continue other medications to treat their IIH (e.g. acetazolamide, diuretics) but this dose should remain fixed throughout the study. Acetazolamide may be taken but the participant must be on a stable dose. |
| 6. | Must be able to understand the consent form and comply with study requirements. |
| Exclusion Criteria | |
| 1. | Optic nerve sheath fenestration. Participants who have had previous failed CSF shunting will be eligible for enrolment if the shunt has failed and they fulfil all other enrolment criteria. |
| 2. | Abnormal neurological examination (aside from papilledema and consequent visual loss or VI nerve palsy). |
| 3 | Unable to perform a visual field reliably. |
| 4. | Positive urine dipstick pregnancy test or planning to conceive in the 4 study months or breast feeding. |
| 5. | Have estimated Glomerular Filtration Rate (eGFR) calculated by Modification of Diet in Renal Disease equation of <60ml/min/1.73m^2^. |
| 6. | Have any complex endocrine disorder, e.g. thyroid dysfunction. Except those with polycystic ovary syndrome and diabetes can be included in the trial. |
| 7. | Suspicion of or known Gilbert's disease. |
| 8. | Creatine kinase >2 x upper limit of normal on 2 consecutive measurements. |
| 9. | Alanine Transaminase and/or aspartate transaminase >2 x upper limit of normal. |
| 10. | Alkaline phosphatase > upper limit of normal. |
| 11. | Bilirubin (total) > 2 x upper limit of normal. |
| 12. | Must not have donated blood within 2 months of screening and avoid further donation for 4 months following the study. |
| 13. | Participant is, at the time of signing the informed consent, a user of recreational or illicit drugs (including marijuana) or has had a recent history (within the last year) of drug or alcohol abuse or dependence, in the opinion of the investigator. |
| 15. | Have uncontrolled systemic hypertension (blood pressure >160 systolic on 3 successive measurements on the screening visit. |
| 16. | Are receiving systemic (including vaginal/rectal) glucocorticoid treatment at the time of the screening visit. |
| 17. | Are taking any hormone-based medication, including hormone contraceptives (but not including IUS/hormonal coil), at the time of screening. |
| 18. | Are taking probenecid at the time of the screening visit. |
| 19. | Have any screening laboratory abnormality that, in the investigator’s judgement, is considered to be clinically significant or any screening laboratory value which is outside reference ranges at screening; testing may be repeated once to see if the value returns to within the range, but any laboratory abnormality must be resolved prior to the baseline visit. |
| 20. | History of any clinically significant disease or disorder which, in the opinion of the investigator, may either put the subject at risk because of participation in the study, or influence the results or the subject’s ability to participate in the study. Specifically, a diagnosis of any inflammatory disorder that might reasonably need treatment with glucocorticoids during the course of the study should be considered for exclusion. |
| 21. | History or presence of significant gastrointestinal, hepatic*, or renal disease or any other condition known to interfere with absorption, distribution, metabolism, or excretion of drugs. [* a history of steatosis will not be considered an exclusion criterion] |
| 22. | Any clinically significant illness, medical/surgical procedure or trauma within 4 weeks of the first administration of the Investigational Medicinal Produce as judged by the investigator. |
| 23. | Have been involved in the planning and/or conduct if the study. |
| 24. | Have participated in any other interventional studies within 1 month prior to the screening visit. Participation in the IIH:Life National database or other observational studies will not prevent enrolment to this study. |
| 25. | Previous randomisation for treatment in the present study. |

**Table 2 supplemental: IIH symptoms**

| **IIH Symptoms** | | **Risk Ratio**  **(95% CI)** | ***p* value** |
| --- | --- | --- | --- |
| **Headaches** | Week 12 | 0.92 (0.64, 1.32) | 0.6 |
|  | Week 16 | 0.78 (0.50, 1.20) | 0.3 |
| **Subjective visual loss** | Week 12 | 0.66 (0.28, 1.57) | 0.3 |
|  | Week 16 | 1.12 (0.47, 2.66) | 0.8 |
| **Pulsatile tinnitus** | Week 12 | 1.06 (0.58, 1.95) | 0.8 |
|  | Week 16 | 0.84 (0.47, 1.52) | 0.6 |
| **Diplopia** | Week 12 | 1.41 (0.14, 13.86) | 0.8 |
|  | Week 16 | 0.71 (0.05, 10.21) | 0.8 |
| **Transient visual obscurations** | Week 12 | 0.83 (0.18, 3.96) | 0.8 |
|  | Week 16 | 0.28 (0.03, 2.22) | 0.2 |

Patient reported occurrence of symptoms (yes / no).

A risk ratio less than 1 favours AZD4017.

**Table 3 supplemental: Visual function and optic nerve head at baseline and week 16**

|  | **Baseline**  **Mean value (SD)** | | **Week 16**  **Mean value (SD)** | | **Adjusted mean difference at 16 weeks (95% C.I)** | **P value** |
| --- | --- | --- | --- | --- | --- | --- |
| **Worse Eye** | **Placebo** | **AZD4017** | **Placebo** | **AZD4017** |  |  |
| **Visual Acuity LogMAR** | 0.13 (0.22) | 0.08  (0.23) | 0.01  (0.26) | 0.04  (0.18) | 0.05  (-0.11, 0.20) | 0.5 |
| **Contrast Sensitivity** | 1.63 (0.16) | 1.63  (0.22) | 1.68  (0.12) | 1.58  (0.44) | 0.02  (-0.09, 0.12) | 0.7 |
| **Perimetric Mean Deviation** | -3.4 (6.8) | -6.1  (5.4) | -1.8  (3.9) | -3.6  (3.5) | 0.6  (-2.0, 3.3) | 0.6 |
| **OCT RNFL Average (μm)** | 158.4 (83.0) | 152.0  (68.7) | 150  (98.2) | 130.0  (46.5) | -17.5  (-54.8, 19.8) | 0.3 |
| **OCT Maximal RNFL (μm)** | 290.0 (102.4) | 320.2  (117.2) | 274.8  (129.0) | 274.1  (98.7) | -27.2  (-95.7, 41.3) | 0.4 |
| **Average Frisén grading** | 2.27 (0.90) | 2.19  (1.17) | 2.17  (0.94) | 1.42  (1.08) | -0.8  (-1.6, 0.02) | 0.06 |

All measures shown in the table are of worst eye. Negative values in the adjusted mean difference between treatment arms favour AZD4017. C.I: confidence interval; OCT: optical coherence tomography; RNFL: retinal nerve fibre layer; SD: standard deviation.

**Table 4 supplemental: Headache outcomes**

|  | Baseline/Screening  Mean values (SD) | | Week 12  Mean values (SD) | |  |  | Week 16  Mean values (SD) | | |  |  |
| --- | --- | --- | --- | --- | --- | --- | --- | --- | --- | --- | --- |
|  | Placebo | AZD4017 | Placebo | AZD4017 | Adjusted mean difference at 12 weeks (95% CI) | p-value | | Placebo | AZD4017 | Adjusted mean difference at 16 weeks (95% CI) | p-value |
| HIT-6 | 63.4 (8.1) | 63.8 (8.2) | 59.8 (7.9) | 61.0 (11.6) | -1.0 (-8.2, 6.1) | 0.8 | | 58.3 (10.6) | 57.1 (8.6) | -0.9 (-7.5, 5.8) | 0.8 |
| Severity  (score 0-10)* | 5.6 (1.8) | 4.6 (1.9) | 4.1 (2.3) | 4.2 (2.8) | 0.4 (-2.0, 2.7) | 0.8 | | 4.2 (2.6) | 3.6 (3.3) | -1.5 (-4.3, 1.3) | 0.3 |
| Duration in 24 hours | 7.6 (4.4) | 7.7 (5.0) | 5.5 (4.3) | 6.8 (6.2) | 1.3 (-3.0, 5.5) | 0.5 | | 5.7 (4.8) | 5.2 (5.7) | -1.5 (-5.8, 2.9) | 0.5 |
| Frequency (days per month) | 22.8 (7.9) | 22.0 (8.6) | 18.2 (11.9) | 17.7 (11.1) | -0.6 (-8.3, 7.1) | 0.9 | | 20.0 (11.8) | 17.6 (12.2) | -2.5 (-11.0, 5.9) | 0.5 |
| Analgesic Use (days per month) | 10.8 (9.6) | 9.6 (9.4) | 5.5 (7.4) | 7.4 (8.9) | 2.6 (-1.6, 6.9) | 0.2 | | 6.8 (11.2) | 2.0 (3.9) | -5.9 (-12.4, 0.6) | 0.07 |

A negative difference favours AZD4017. Values are adjusted from baseline.

*Patient rating scale where 0=no headache to 10=worst pain ever experienced

**Table 5 supplemental: Adverse events by system class**

|  | **Placebo**  **(n=14)** | **AZD4017**  **(n=17)** |
| --- | --- | --- |
| Cardiovascular | 1 (7%) | 1 (6%) |
| Respiratory | 3 (21%) | 4 (24%) |
| Gastro-intestinal | 3 (21%) | 8 (47%) |
| Genito-urinary | 0 (0%) | 6 (35%) |
| Endocrine | 1 (7%) | 1 (6%) |
| Musculoskeletal | 7 (50%) | 6 (35%) |
| Neoplasia | 1 (7%) | 0 (0%) |
| Neurological | 3 (21%) | 7 (41%) |
| Psychological | 4 (29%) | 4 (24%) |
| Immunological | 1 (7%) | 0 (0%) |
| Dermatological | 5 (36%) | 2 (12%) |
| Allergies | 0 (0%) | 1 (6%) |
| Eyes, ears, nose, throat | 6 (43%) | 12 (71%) |
| Other ^*^ | 5 (36%) | 7 (41%) |

Other complaints included: tiredness; hot sweats; flu like symptoms; disrupted sleep; toothache/infection; breast pain; menstrual problems for more than 3 weeks; mouth ulcers; a cold; transient nausea and height headaches. Data are n (%).

**Table 6 supplemental: Safety blood tests**

|  | **Baseline**  **Mean value (SD)** | | **Mean difference 95% CI** | **12 Weeks**  **Mean (SD)** | | | **Mean difference 95% CI** |
| --- | --- | --- | --- | --- | --- | --- | --- |
|  | **Placebo** | **AZD4017** |  | **Placebo** | | **AZD4017** |  |
| **Renal Function (mmol/L)** |  |  |  |  | |  |  |
| Urea | 3.57 (0.93) | 4.04 (1.39) | 0.47  (-0.44, 1.39) | 3.64 (1.04) | | 3.65 (0.99) | 0.01  (-0.78, 0.79) |
| Creatinine | 66.54 (8.00) | 65.18 (8.10) | -1.36  (-7.44, 4.72) | 68.17 (8.79) | | 71.24 (11.17) | 3.07  (-4.87, 11.01) |
| Potassium | 4.08 (0.32) | 4.11 (0.22) | 0.02  (-0.19, 0.23) | 4.09 (0.18) | | 4.14 (0.27) | 0.04  (-0.14, 0.23) |
| Sodium | 139.8 (3.61) | 140.1 (1.93) | 0.35  (-1.75, 2.44) | 140.4 (3.00) | | 154.4 (67.02) | 14.00  (-25.9, 53.93) |
| **Liver Function** |  |  |  |  | |  |  |
| Aspartate aminotransferase (AST) (U/L) | 20.85 (10.29) | 20.50 (9.04) | -0.35  (-7.71, 7.02) | 22.56 (6.98) | | 21.59 (5.62) | -0.97  (-6.17, 4.23) |
| Alanine aminotransferase (ALT) (U/L) | 25.62 (11.98) | 25.00 (12.60) | -0.62  (-10.1, 8.83) | 28.08 (16.09) | | 23.29 (9.62) | -4.79  (-14.6, 5.01) |
| Alkaline phosphatase (ALP) (U/L) | 70.31 (17.93) | 62.63 (17.67) | -7.68  (-21.3, 5.94) | 71.00 (18.55) | | 49.82 (13.27) | -21.2  (-33.3, -9.08) |
| Bilirubin (μmol/L) | 5.46 (1.94) | 6.59 (2.87) | 1.13  (-0.77, 3.03) | 5.83 (2.48) | | 6.18 (1.63) | 0.34  (-1.22, 1.91) |
| Albumin (g/L) | 42.80 (2.17) | 41.25 (1.26) | -1.55  (-4.46, 1.36) | 43.00 (2.00) | | 44.00 (1.83) | 1.00  (-2.06, 4.06) |
| γ-glutamyltransferase (U/L) | 30.77 (14.04) | 24.31 (13.75) | -6.46  (-17.1, 4.18) | 27.33 (13.01) | | 17.24 (8.45) | -10.1  (-18.3, -1.94) |
| **Thyroid Function** |  |  |  |  | |  |  |
| Thyroid stimulating hormone (mlU/L) | 1.41 (0.68) | 1.76 (0.55) | 0.35  (-0.11, 0.80) | 1.71 (0.65) | | 1.87 (0.46) | 0.16  (-0.26, 0.59) |
| Free Thyroxine (pmol/L) | 16.65 (8.59) | 14.68 (2.55) | -1.97  (-6.46, 2.51) | 13.56 (1.43) | | 14.82 (2.04) | 1.26  (-0.15, 2.66) |
| **Muscle Function** |  |  |  |  | |  |  |
| Creatine Kinase (U/L) | 93.75 (38.99) | 106.2 (78.46) | 12.43  (-36.6, 61.46) | 107.7 (48.54) | | 97.29 (49.83) | -10.4  (-48.5, 27.77) |
| **Hypothalamic Pituitary Adrenal Function** | | | | | | | |
| Cortisol (nmol/L) | 245.6 (89.85) | 243.7 (118.5) | -2.00  (-80.7, 76.69) | 240.0 (63.16) | 214.5 (112.2) | | -25.5  (-106, 55.00) |
| Dehydroepiandrosterone Sulphate | 6.29 (2.91) | 6.92 (3.33) | 0.63  (-1.82, 3.08) | 5.54 (3.39) | 10.98 (6.39) | | 5.44  (1.09, 9.79) |
| 4-Androstenedione (nmol/L) | 4.78 (2.90) | 4.69 (2.14) | -0.08  (-1.97, 1.80) | 5.22 (2.87) | 5.53  (1.98) | | 0.31  (-1.57, 2.20) |
| Testosterone (nmol/L) | 1.65 (1.25) | 1.35 (0.64) | -0.30  (-1.02, 0.42) | 1.61 (0.70) | 3.00  (4.87) | | 1.39  (-1.53, 4.32) |
| Adrenocorticotropic Hormone (ng/L) | 17.13 (11.29) | 14.01 (10.28) | -3.11  (-11.5, 5.30) | 13.02 (10.87) | 25.38 (18.57) | | 12.36  (-0.03, 24.74) |
| Follicle Stimulating Hormone (IU/L) | 12.96 (22.10) | 5.90 (2.84) | -7.06  (-18.1, 3.98) | 13.00 (13.49) | 6.59  (4.27) | | -6.41  (-13.5, 0.72) |
| Luteinising Hormone (IU/L) | 10.49 (15.14) | 6.82 (4.36) | -3.67  (-11.6, 4.21) | 11.96 (8.35) | 8.19  (6.01) | | -3.77  (-9.23, 1.69) |
| Oestrodiol (pmol/L) | 309.8 (247.5) | 336.0 (226.0) | 26.25  (-165.0, 218.0) | 240.9 (182.9) | 312.9 (213.3) | | 71.97  (-108, 251.7) |
| Progesterone (nmol/L) | 3.25 (3.88) | 7.34 (12.73) | 4.08  (-4.17, 12.34) | 1.93 (1.52) | 10.89 (17.44) | | 8.96  (-2.59, 20.50) |

**Table 7 supplemental: AZD4017 drug concentrations in serum and CSF at baseline, week 1 and week 12.**

| Week | Mean concentration (ng/ml)±SD | |
| --- | --- | --- |
|  | Serum (n=6) | CSF (n=6) |
| 0 | 0±0 | 0±0 |
| 1 | 1861±1067 | N/A |
| 12 | 1957±1874 | 8.95±2.87 |
